# Production of photocurrent and hydrogen gas from intact plant leaves

**DOI:** 10.1101/2021.09.19.460952

**Authors:** Yaniv Shlosberg, Matan Meirovich, Omer Yehezkeli, Gadi Schuster, Noam Adir

## Abstract

Here, we show that it is possible to harvest photocurrent directly from unprocessed plant tissues from terrestrial or aquatic environments in bio-photoelectrochemical cells (BPECs) and use the current to produce molecular H_2_. The source of electrons is shown to originate from the Photosystem II water-oxidation reaction and utilizes exported mediating molecules, especially NADPH. The photocurrent production is dependent on the concentration of the photosynthetic complexes, as an increase in total chlorophyll and oxygen evolution rates lead to increased photocurrent rates. The permeability of the outer leaf surface is another important factor in photocurrent harvesting. Different tissues produce photocurrent densities in the range of ∼ 1 – 10 mA / cm^2^ which is significantly higher than microorganism-based BPECs. The relatively high photocurrent and the simplicity of the plants BPEC may pave the way toward the development of future applicative photosynthetic based energy technologies.

## 1. Introduction

World energy demands continue to increase steeply due to population growth and lifestyle requirements. It was estimated that by 2035, global energy consumption will reach an annual average of ∼26 TW.(Khatib, 2012) Current fossil fuel combustion technologies handle most energy generation requirements, however these lead to greenhouse gas emission and which may be a source of climate change (Ritchie et al., 2021; Schleussner et al., 2016). The increasing awareness of fossil fuel pollution has led to the search for cleaner renewable energy technologies. These include utilization of biological systems, including bioelectricity generation technologies which are based on the harvesting of electrical current produced by live organisms (Fischer, 2018; González del Campo et al., 2014; Kabutey et al., 2019; Rosenbaum et al., 2010; Strik et al., 2008) or by utilizing isolated enzymes (Hartmann et al., 2020; Herzallh et al., 2020; Nam et al., 2018) for the generation of light-induced electrical power.

The first report of use of microbes for the generation of electrical energy by microbial fuel cells (MFCs) was made by Potter et al. in 1910 (Ieropoulos et al., 2005). Establishment of efficient electrical communication between bacteria and electrodes can be obtained by either direct electron transfer (DET) (Lovley, 2012; Nevin et al., 2008; Yi et al., 2009) or mediated electron transfer (MET) (Chaudhuri and Lovley, 2003; Liu et al., 2006; Min et al., 2005; Pham et al., 2003). DET has been reported for the bacterial species such as *Geobacter sulfurreducens* and *Shewanella oneidensis* that use conductive inter-membranal protein complexes such as found in *pili* or metal respiratory system (MTR) complexes to export electrons (Hartshorne et al., 2007; Shi et al., 2012). MET systems are more prevalent as they can be performed either by addition of exogenous mediators or by the secretion of endogenous reducing molecules by the cells (Bond and Lovley, 2003; Chaudhuri and Lovley, 2003; Liu et al., 2006; Min et al., 2005; Pham et al., 2003). To achieve enhanced electrical current generation in MFCs, artificial electron mediators such as potassium ferricyanide (FeCN), and various quinones or phenazines derivatives have been added (Ieropoulos et al., 2005; Lovley et al., 1996; Rabaey et al., 2005, 2004; Simoska et al., 2019). MFCs typically require a constant flow of solutions of organic molecules of specific composition optimized for the organism’s long-term survival and electron transfer activity.

A different strategy for bio-electrical power generation, is based on photosynthetic organisms. Intact photosynthetic microorganisms, organelles such as chloroplasts, (Hasan et al., 2017) thylakoid membranes (Pinhassi et al., 2016) or isolated photosynthetic complexes such as Photosystem II (PSII) (Hartmann et al., 2020; Riedel et al., 2019; Shoyhet et al., 2021; Yehezkeli et al., 2012) or Photosystem I (PSI) (Efrati et al., 2016, 2013; Herzallh et al., 2020; King, 2018; de Moura Torquato and Grattieri, 2022; Yehezkeli et al., 2012; Zhao et al., 2014) have been coupled with electrodes in different bio-photo electrochemical cells configurations. Upon illumination, these photosynthetic components convert absorbed light energy into electrical current collected by the anode. Similar to some MET type MFCs, the electron transfer from these photosynthetic components to the anode may be enabled or enhanced by the addition of an exogenous mediators such as potassium ferricyanide or quinones or via tightly bound redox polymers or nanoparticles (Weliwatte et al., 2021). Although isolated photosystems can theoretically provide a high concentration of photochemical reaction centers (and thus high current densities), their isolation is more complicated and expensive than the use of isolated photosynthetic membranes or intact organisms and therefore may be less attractive for practical applications.

While the direct use of electrical current harvested from photosynthesis may be an attractive alternative energy source, use of the current to produce an energy-rich compound such as H_2_ has advantages as it can be stored and used in existing technologies. Indeed, recent research has shown the potential of bio-photo electrochemical cells which are based on photosynthetic components to produce molecular H_2_ (King, 2018; Yehezkeli et al., 2012), that can be utilized for pollution-free electricity production in hydrogen-based fuel cells (Robledo et al., 2018). H_2_ evolution may be achieved by utilizing platinum cathodes to catalyse the reduction of H^+^ into H_2_ (Pinhassi et al., 2016) or by utilizing the activity of enzymes such as hydrogenase or nitrogenase that produce H_2_ (Herkendell et al., 2017; Milton et al., 2017, 2016; Yehezkeli et al., 2012).

BPECs based on intact, live photosynthetic microorganisms such as cyanobacteria or microalgae can transfer photoexcited electrons to the anode to generate electrical current in a fashion similar to non-photosynthetic organism based MFCs (Lai et al., 2021; Li et al., 2021; Saper et al., 2018; Shlosberg et al., 2020; Wang et al., 2014). These photosynthetic microorganisms can utilize atmospheric CO_2_ to synthesize their carbon source and therefore, are not dependent on a constant external source of carbon. (Tomar et al., 2017) Moreover, species such as *Dunalliela* (Borovkov et al., 2020; Hosseini Tafreshi and Shariati, 2009) or *Arthrospira platensis* (*Spirulina*) (Silva et al., 2020) are already cultivated in industrial facilities around the world for the production of food additives and other high value products. These well-established technologies are an excellent platform to integrate bioelectricity or hydrogen production into existing cultivation facilities. Recently it has been shown that the source of the external electron transport in cyanobacteria derives from a combination of the respiratory and photosynthetic pathways. (Saper et al., 2018) Unlike in non-photosynthetic exo-electrogenic bacteria, it was shown that type IV *pili* in cyanobacteria, is not involved in the external electron transfer (EET) mechanism (Thirumurthy et al., 2020). It was further shown that both cyanobacterial and unicellular green algae cells, in association with a BPEC anode, secrete Nicotinamide adenine dinucleotide phosphate (NADPH) which acts as the major electron mediator (Hatano et al., 2022; Shlosberg et al., 2021c, 2020). Upon illumination, more NADPH is formed by the photosynthesis pathway, leading to an enhanced secretion process which subsequently leads to higher currents. The resulting NADP^+^ can be recycled back into the cells, to be re-reduced by photosynthetic ferredoxin-NADP^+^ reductase. Furthermore, the addition of exogenous NADP^+^ (or NAD^+^) was reported to significantly enhance the photocurrent intensity and duration (Shlosberg et al., 2020). Using a different approach, enhanced photocurrents were gained by cyanobacterial biofilms grown directly on an anode surface (Gonzalez-Aravena et al., 2018; Ng et al., 2017).

We have recently shown that bio-photo electrochemical cells are not limited to microorganisms, and that macroalgae (seaweeds) can be utilized for bioelectricity production as well (Shlosberg et al., 2021b). As the macroalgae thrive in an aqueous environment, the photosynthetic tissues import all nutrients through the outer thallus surface as well as secrete molecules to control the immediate surrounding. A series of studies have previously revealed that such a potential benefit for connection to a BPEC can also be obtained with aquatic higher plants such as *Lemna minuta* (Duckweed) (Hubenova et al., 2018). These studies showed that these plants can be grown in direct connection with carbon-felt based electrodes, providing light-dependent current over many days. As the photosynthetic apparatus in all plants are sequestered within chloroplasts, these observations indicate that electrons can be transported out of the chloroplast and out of the cells and leaf cuticle. No similar study has been reported for non-aquatic plants, which have a variety of outer surfaces, and are typically exposed to air, and not water. However, if electrons could be harvested in a similar fashion from terrestrial plants, the potential of such technologies would potentially expand to almost any terrestrial environment. Terrestrial plants have a wide range of tissues that are appendages of a main body (typically the stem) which terminates with a root system attaching the plant to the ground or other stable surface. These tissues have evolved to perform three major physiological tasks – light absorption for photosynthesis, water absorption and gas exchange. These different functions must work in concert and under the proper control to maximize solar energy conversion to storable sugars, while also providing protection: from photoinhibition (Adir et al., 2003; Pinnola and Bassi, 2018), desiccation (Bechtold, 2018) and from other forms of biotic or abiotic stress (Li et al., 2017). Synchronization of these different functions requires the movement of various metabolites throughout the tissues (leaves, fronds, needles, microphylls, sheaths, etc.; the general term “leaves” will be used hence), crossing various cell membranes into the extracellular spaces, and within the cells into the chloroplasts and mitochondria. While aqueous plants environments absorb carbon from dissolved CO_2_, terrestrial plants absorb CO_2_ and water through dedicated stomata which can open or close (Lawson and Matthews, 2020). While it might seem counterintuitive to suggest that it is possible to harvest electrons (via reduced molecules) from leaves, the concentrations of such molecules are not trivial (Rodrigues and Shan, 2021). Here, we show that terrestrial plants can generate photocurrents in BPECs using different tissues such as leaves and stems. Finally, we demonstrate the concept of using aqueous plants to generate electrical current using their native environment under natural sunlight irradiation.

## 2. Materials and Methods

### Materials

All chemicals were purchased from Merck. ***Spinacia oleracea*** leaves were purchased at local markets in Haifa.

### Indoor CA measurements leaves

The indoor CA measurements of flat leaves were done in the same setup which was described in our previous work (Shlosberg et al., 2021b). The measurements of all leaves were done in a small rectangular transparent glass vessel with dimensions of 4.5 cm^3^. A solar simulator (Abet, AM1.5G) was placed horizontally to illuminate the leaves with a solar intensity of 1 Sun (1000 W/m^2^). Determination of the light intensity at the surface of the leaves was done as a function of distance from the light source in an empty vessel neglecting small intensity losses caused by the glass and (∼ 0.5 cm) of the electrolyte solution. The measurements were conducted in 3 electrode mode (unless otherwise mentioned) using the stainless-steel clip as anode, a platinum wire as a cathode, and Ag/AgCl 3M NaCl as a reference electrode (RE-1B, CH Instruments, USA) with an applied electric potential bias of 0.5 V on the anode in NaCl solution (0.5 M). In all measurements, the current density was calculated by subtraction of the stable baseline after ∼ 10 min and based on the contact area between the anode and the leaves of 0.08 cm^2^. When applied, the addition of 3-(3,4-dichlorophenyl)-1,1-dimethylurea (DCMU) was done prior to the measurements (5 min).

### Hydrogen production and quantification

The BPEC was covered by a thick parafilm layer. The volume of the electrolyte solution was 50 mL and the headspace volume was 40 mL. Following 10 min of CA measurements, 1 mL of air was removed from the top of the reaction vessel and injected into vials (1.8 mL). samples (50 μL) were injected into a gas chromatograph (GC) system coupled with a thermal conductivity detector (GC-TCD, Agilent 8860) with a 5-Å column (Agilent, 25m x 0.25mm x 30μm). Hydrogen that evolved during the BPEC stabilization stage (see previous section) were subtracted from the values of hydrogen obtained during the actual experiment.

### Spectroscopic fluorescence and microscopic measurements of spinach and its external solution

Absorption measurements of the external solution of spinach was done by Shimadzu (UV-1800) spectrophotometer in 1 cm pathlength square cuvettes. 2D - fluorescence measurements external solution of spinach were performed using a Fluorolog 3 fluorimeter (Horiba) with excitation and emission slits bands of 4 nm as previously described(Shlosberg et al., 2020).2D - fluorescence measurements from the spinach leaf were done directly from the surface of the leaf that was placed with an angle of 45 degrees between the illumination source and the detector of the fluorimeter. The lines of diagonal spots that appear in all maps result from the light scattering of the Xenon lamp and Raman scattering of the water. Fluorescence intensity is presented as calibrated color maps in units of counts per second (CPS).

Spinach cuttings were grasped by the anode and incubated for 20 min in 0.5 M NaCl. The grasped area of the spinach was observed and photographed by Horiba Jobin Yvon (LabRAM HR Evolution®) Micro-Raman.

### Direct CA measurements from the water lily pond

CA measurements were done directly from the pools using a stainless-steel clip anode, a Pt wire as a cathode, and Ag/AgCl 3M NaCl as a reference electrode. The anode clip was grasping a waterlily leaf. The anode and reference electrode were inserted into a sponge that was floating on the pool surface. An applied electric potential of 0.5 V on the anode under the sunlight. Light intensity was measured at the water surface height of the pond with an app-based portable light meter Lux The temperature was monitored manually during the duration of the measurement.

### Dissolved oxygen measurements

Dissolved oxygen (DO) measurements were conducted in the same system of the CA measurements with the addition of a small magnetic stirrer bar. The top of the reaction vessel was tightly covered with a thick layer of parafilm. A DO meter probe (Hanna Instruments, HI-5421 research grade DO and Biochemical oxygen demand bench meter) was inserted into the BPEC liquid phase (50 ml) to quantify the accumulated DO concentration after 10 min.

### Chlorophyll concentration determination

Chlorophyll determination was done by grinding of the leaves followed by acetone extraction followed by absorption measurements as previously described (Ritchie, 2008, 2006)

### Determination of the pressure applied to spinach leaves by the anode clips

The pressure imparted by the stainless steel clips that serve as the anode on the leaf tissues were measured with a universal load testing machine (MTS), in the Faculty of Aerospace Engineering, Technion.

### 3. Results and Discussion

#### 3.1 *Spinacia oleracea* (Spinach) produces photocurrent and hydrogen in a BPEC using electrons that originates from Photosystem II

In our previous studies on the design of cyanobacterial (Saper et al., 2018; Shlosberg et al., 2020) and green algae BPECs (Shlosberg et al., 2021c) the association between the cells and the anode could be conducted simply by placing the cells (in the appropriate electrolyte solution) directly on the anode. Recently, we have shown that photocurrents can also be harvested from macroalgae that have large leaf-like (thallus) tissue that must be held in place to physically contact electrodes (Shlosberg et al., 2021b). This alternative BPEC setup was found to be useful for terrestrial leaves as well. This BPEC consists of a transparent glass container, stainless steel (Type 304) flat anode clips (without sharp metallic teeth, Fig. S1), that holds the leaf, a platinum cathode wire and an Ag/AgCl 3M NaCl as the reference electrode (Fig. 1a and S2). In our previous work with seaweeds based BPECs (Shlosberg et al., 2022), we showed that the performance of stainless steel is better than other materials such as aluminium, carbon cloth, or indium tin-oxide. The reason for this may derive from the high conductivity of stainless steel, but also from a better biocompatibility. A main component of stainless steel is iron that in nature can be reduced by plant’s roots (Bienfait, 1985) or leaf extracts (Mehrotra and Gupta, 1990). We speculated that plants leaves can also perform iron reduction which might be the reason for the better biocompatibility of stainless steel over other kinds of electrodes. Over time stainless-steel does slowly corrode. This may be a disadvantage that will require the replacement of the electrodes once they become highly damaged. However, the cost of simple electrodes such as stainless is relatively low. Chronoamperometry (CA) measurements were performed by directly connecting round segments of leaf from spinach (purchased at local markets) with a diameter of ∼1 cm to the stainless-steel clip either in dark or light. The CA measurements were conducted with an applied bias potential of 0.5 V on the anode. This bias value was chosen based on our previous studies of BPECs based on different photosynthetic organisms, that showed that this is an optimal value for maximal photocurrent production (Shlosberg et al., 2021b, 2021c, 2020). The application of this potential bias was previously reported to interact with photosynthetic cells membrane enhancing the release of the electron mediator NADPH (Shlosberg et al., 2020).To optimize the electrolyte concentration, CA measurements were conducted in electrolyte solutions with NaCl concentrations between 1-750 mM (Fig. S3), with photocurrents increasing up to 500 mM NaCl (Fig. S3). Prior to every measurement, the measured current in the BPEC system was allowed to stabilize in the dark which takes 10 min. When the leaf cutting is illuminated, the BPEC produced maximal photocurrents of 6.7 +/- 0.4 mA / cm^2^ after 10 min (Fig. 1b, c). This photocurrent production is significantly higher than previously obtained from photosynthetic microorganisms (Shlosberg et al., 2021c, 2020). We postulated that the major reasons for this are the density of photosynthetic membranes (that is significantly higher in plants leaves as opposed to cyanobacteria or green algae) coupled to the ability to tightly grasp a leaf, strengthening the functional MET connection with the anode. The measured open circuit potential of the BPEC was 0.37 V. Solar efficiency is the ratio between the maximal power output and the solar irradiation intensity (1000 W / m^2^). The calculated solar efficiency was fond to be 2.51 +/- 0.15 %, for the area of leaf active in electron transfer to the anode (see below). This efficiency is about 10 % of what has been estimated to be typical photosynthetic efficiency starting from PSII water oxidation (Blankenship, 2001). This value is about 10% of the quoted efficiency of most commercial solar cells (Day et al., 2019). To further explore the kinetics of dark and light reaction, CA was measured with cycles of 10 min in light followed by 5 min in dark (Fig. S4). The results showed that the increase in the photocurrent, in the first light regime, was slower than its decrease when the light was switched off. However, in the following light/dark periods, the rate of increase/decrease in the current were similar. This suggests that during the original association period, the operation of the BPEC influences the release of mediating molecules, that is then preserved over time. To evaluate the sustainability of the photocurrent over a longer period of time, CA of spinach was measured for 1 h in light showing that a maximal photocurrent density of ∼6.5 mA /cm^2^ was sustained (Fig. S5). This observation shows the benefit of using live organisms, as when membranes or isolated photosynthetic components are utilized in BPECs, the photodamage to the reaction centres (especially PSII) limits the lifetime of the photosynthetic system due to photoinhibition that cannot be repaired (Adir et al., 2003; Li et al., 2018).

**Fig. 1.**
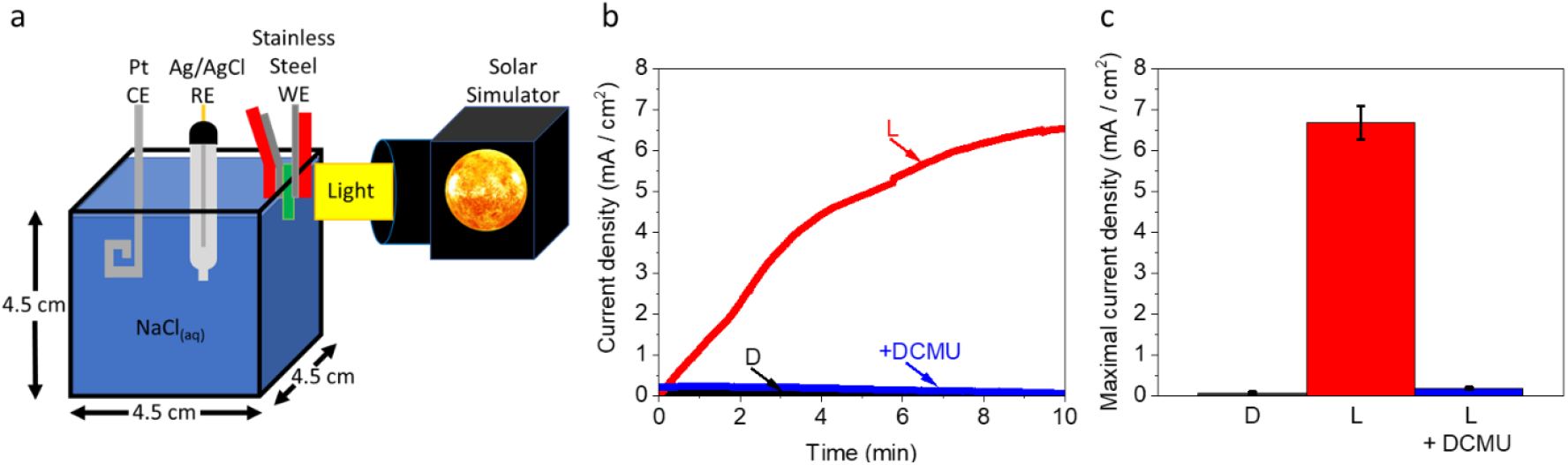
Intact spinach leaves produce photocurrent from Photosystem II in a BPEC. **a**. CA measured using a leaf disk. A maximal photocurrent of ∼ 6.7 +/- 0.4 mA / cm^2^ was obtained when the leaf disk was illuminated with white light (red). No significant current was obtained in either dark (D, black) or in the presence of DCMU (blue). **b**. Maximal current production in the CA of spinach leaves in dark (D, black), light (L, red), and light + DCMU (blue). The error bars represent the standard deviation over 3 independent measurements.

To measure the hydrogen evolution, the top of the reaction vessel was sealed with a thick layer of parafilm, and CA measurements were performed for 10 min in dark or light, with an applied potential of 0.5 V (vs. Ag/AgCl) on the anode in the absence or presence of 100 μM DCMU. Samples of the head space of the BPEC were removed with an air-tight syringe and the amount of hydrogen was quantified by gas chromatography analysis. We found that the system evolved 4.19 +/- 0.53 μmol hydrogen (average and S.D. of three independent measurements) in the absence of DCMU. No hydrogen was detected in the absence of light or in the light with DCMU present. See Methods for additional details on the hydrogen evolution measurements.

While using a metal electrode in a BPEC, some of the electrical current may derive from corrosion of the metal. The current density that was obtained for CA measurements of spinach in dark and in light with the addition of DCMU was negligible (Fig.1), indicating that the contribution of corrosion to the photocurrent is minute. To further evaluate the possible contribution of corrosion to current generation, CA was measured in the absence of a spinach cutting, for 10 min with or without addition if 100 μM DCMU. The corrosion-based photocurrents with and without DCMU were estimated to be ∼ 0.05 mA /cm^2^, or less than 1% of the photocurrent obtained in the presence of the spinach cutting (Fig. S6). In this control, the current obtained with or without DCMU was similar, showing that DCMU does not affect corrosion-based current.

Since the electrochemical reaction occurs at the surface of the electrodes, it is common to normalize the electrical current production to the anode surface (current density). However, the current formation in the BPEC was formed only upon irradiation, while the anode clip itself is covered by plastic and thus not exposed to the light while the leaf cutting is. Thus, upon illumination mediated electron transport is driven by reduced molecules released from the cells (and chloroplasts found within). This raised the question whether the optimal normalization should relate to the anode surface area or illuminated leaf area. Illumination is performed on one side of the leaves; however, the light penetrates the leaf interior, and molecules can be released from interior cells as well. We performed CA on spinach cuttings exposed to anode areas of 0.08 and 0.04 cm^2^ (Fig. S7). The measured current ratio was 1.82 showing that the anode surface does indeed affect the amount of current harvested. To evaluate the effective distance in the leaf that influence photocurrent production, spinach leaves were cut so that they had a fixed width of 0.2 cm (the width of the anode) and an exposed length of 0.0 – 0.3 cm (the smallest piece is thus not exposed to light (Fig. S8a, b). The maximal effective distance in the leaf that contributes to photocurrent production was evaluated to be 0.1 cm. (Fig. S8c).

These experiments show the usefulness of using intact leaves as this allows the use of elevated ionic strength buffers as the electrolyte solution. The elevation of the ionic strength has shown to promote orders of magnitude increases in current densities in microbial fuel cells (Huang et al., 2009; Liu et al., 2005) and in photosynthetic organism based BPECs. (Shlosberg et al., 2021b; Tschörtner et al., 2019)

### 3.2 Tight attachment of leaves to the anode enhances the photocurrent generation by enhancing the release of NADPH

In our previous studies (Shlosberg et al., 2021b, 2021c, 2020), we showed that NADPH that is formed in the photosynthetic pathway serves as the major electron mediator that can shuttle electrons between PSI inside the chloroplast and cytoplasm to the anode to produce electrical current. This electron transport mechanism was reported for BPECs based on cyanobacteria, microalgae, and macroalgae. We wished to assess whether a similar mechanism exists also in plants based BPECs. Also, we wished to assess the role of the mild pressure applied to the leaf by the anode clips in the BPEC on the possible increase in the release of NADPH from the leaf. We first measured 2D-fluorescence maps from the surface of an intact spinach leaf (λ_Ex_ = 250 – 500 nm, λ_Em_ = 280 – 700 nm; Fig. 2a). The spectral fingerprints of the amino acids Tyrosine and Tryptophan (Tyr + Trp) (λ_Ex_ max= 280 nm, λ_Em_ max = 350 nm)(Shlosberg et al., 2021a), NADPH (λ_Ex_ max= 280 nm, λ_Em_ max= 350 nm)(Shlosberg et al., 2020), and chlorophyll a (λ_Ex_ max= 480nm, λ_Em_ max = 690 nm)(Shlosberg et al., 2020) were identified. Intact spinach cuttings and cuttings grasped by the non-sharp edge of the anode clips (as in the BPEC setup, Fig. 1a) were incubated for 20 min in 0.5 M NaCl. The spinach cuttings were removed, and absorption spectra of the BPEC solutions were measured. The spectra showed peaks at λ = 250 – 450 nm (Fig. 2b). As many biomolecules absorb in this spectral region, the absorption spectra was not informative enough to identify specific molecules that are released from spinach cuttings. However, the peak of the incubation solution that contained the spinach cutting grasped by the clip anode was ∼ 3 times higher than from the leaf cutting itself. Based on these results, we postulated that the pressure applied by the anode clips, enhances the general release of molecules to the external solution. To identify the release of NADPH from the spinach pieces, 2D-fluorescence spectra of the solutions of the native and anode grasped spinach were measured. Similar to the results obtained by absorption measurements, the spectral fingerprint of NADPH was clearly identified in both solutions, while the intensity of the incubation solution of the grasped leaves was about 3 times greater than the solution of the intact cuttings. (Figs. 2c, d). We concluded that the increased release of NADPH molecules from the leaves by the mild pressure of the anode may be a factor that enhances current production. A similar phenomenon was previously observed in cyanobacterial based BPECs, in which the cells were treated with a mild pressure using a microfluidizer (Saper et al., 2018). The mechanical pressure of the clips imparted by the spring on the spinach cutting was found to be 10 N/mm^2^ (see Methods for additional details). To ascertain whether grasping the leaf by the anode physically damages the leaf tissue, we examined the area of the spinach that was grasped and incubated for 20 min in NaCl solution was observed using an optical microscope. The microscopic photos showed that no physical damage was done to the leaf tissue while the cells wall boundaries are well observed as intact (Fig. S9). Although the grasp of the anode clip did not physically harm the leaf, it may cause various effects over time. The grasp of the leaf can change the turgor pressure influencing the activity of water fluxes depend on plasmodesmata, aquaporins, transporters and channels, and influence developmental regulation pathways (Beauzamy et al., 2014). As indicated above, activation of the potential bias on the working electrode will affect the organization of ions from the electrolyte solution on both the surface of the electrode and the surface of the leaf.

**Fig. 2.**
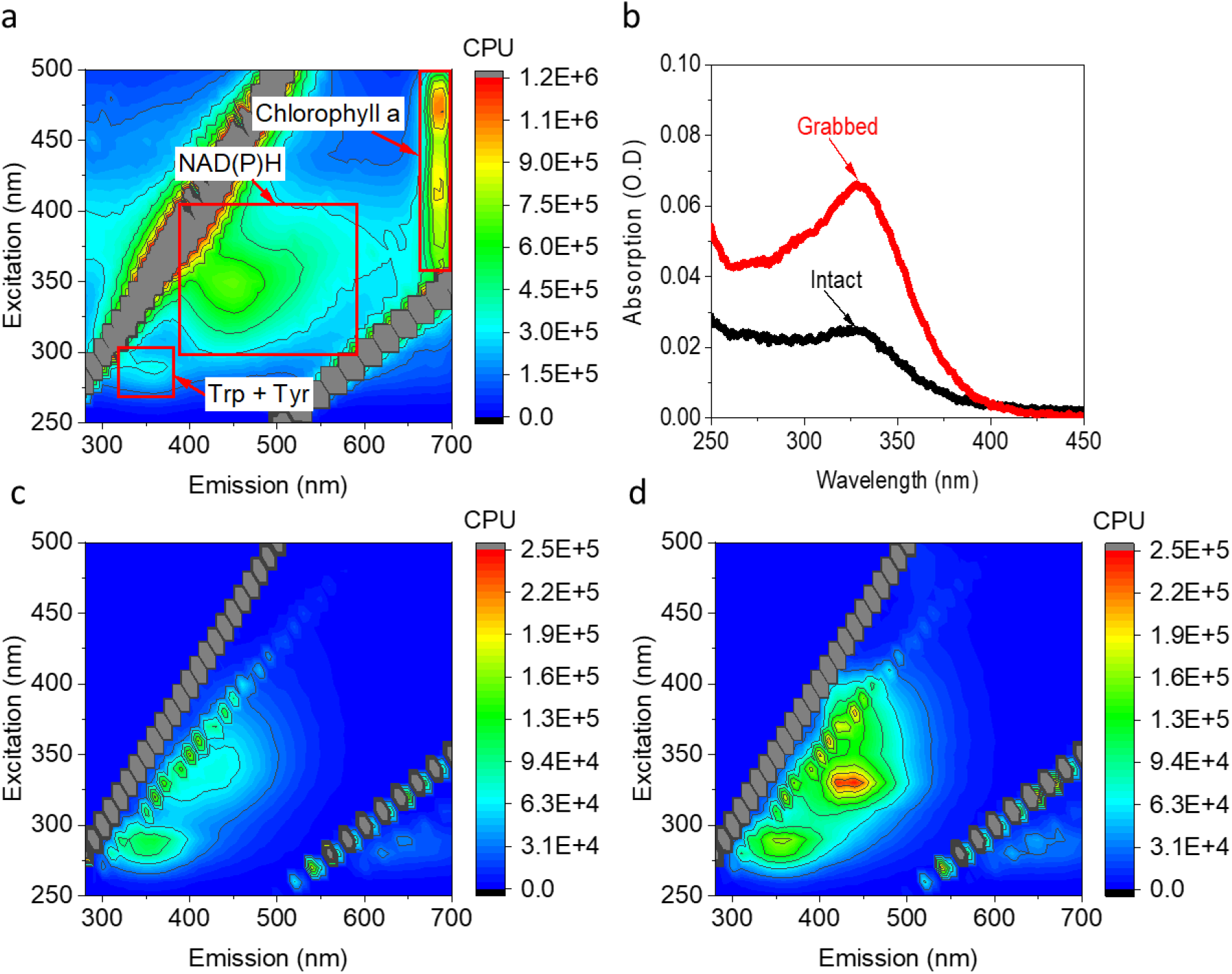
Attachment of leaves to the anode enhances release of NADPH. **a** 2D-fluorescence map of a section of an intact spinach leaf. Red squares mark the peaks of chlorophyll a, NAD(P)H and the amino acids Trp + Tyr **b** Absorption spectra of the electrolyte solution after incubation with intact spinach cuttings (black), or a cutting grasped by the anode clip (red). **c** 2D-fluorescence map of the electrolyte solution after incubation with the intact spinach cutting, not attached to the anode clip. **d** 2D-fluorescence map of the solutions after incubation with the spinach cutting grasped by the anode clip. The lines of diagonal spots that appear in all of the maps presented here and in the following Figs result from the light scattering of the Xenon lamp and Raman scattering of the water (Lawaetz and Stedmon, 2009). The magnitude of fluorescence (counts per second, CPS) is color-coded as indicated on the bar on right of each spectra.

### 3.3 Photocurrent production of plants with different leaf morphologies

The results presented above using spinach in the BPEC designed to hold intact leaves, led us to explore if leaves of plants with other leaf morphologies and growth characteristics, can be coupled to the plant BPEC to generate photocurrent. CA measurements were performed using leaves of various plants from different native habitats. We explored *Salvia officinalis* (Sage), *Pinus* (Pine needles), *Cistaceae* (Cistus), *Salvia Rosmarinus* (Rosemary), *Vitis* (Grapevine), *Bryophyta* (Moss), stems of *Rosa* (Rose) and the *Cacti* plant *Opuntia Ficus-indica*. Our results show that upon illumination, most leaves are able to generate an electric current in the BPEC (Fig. 3), including leaves from trees, bushes, and the microphylls of moss. Planar leaves with soft textures produced ∼ 6 – 9 mA / cm^2^ while photosynthetic tissue (stems or leaves) with hard textures produced lower currents of ∼ 1– 2.5 mA / cm^2^. CA of a dry cistus leaf did not produce any photocurrent (Fig. S10). This validation shows that the leaves must be viable to generate photocurrent. No significant photocurrent could be obtained from intact *Cacti* plant *Opuntia ficus-indica* phylloclade (that consists of short photosynthetically competent stems). However, removal of its external rough layer enabled the harvesting of a photocurrent of ∼ 11 mA / cm^2^. A picture of the *Opuntia ficus-indica* attached to a BPEC, before or after removal of its external layer, is shown in Fig. S11. The results show that the ability of photosynthetic leaves and stems to produce photocurrent is inversely dependent on the toughness of their external cuticle. Based on these results, we suggest that plant leaves that are more permeable (lack a cuticle) enable a higher flux of reducing molecules to exit the cells, while attaching the anode directly to the semi-liquid layer below the *cacti* cuticle provides the highest level of interaction with MET molecules.

**Fig. 3.**
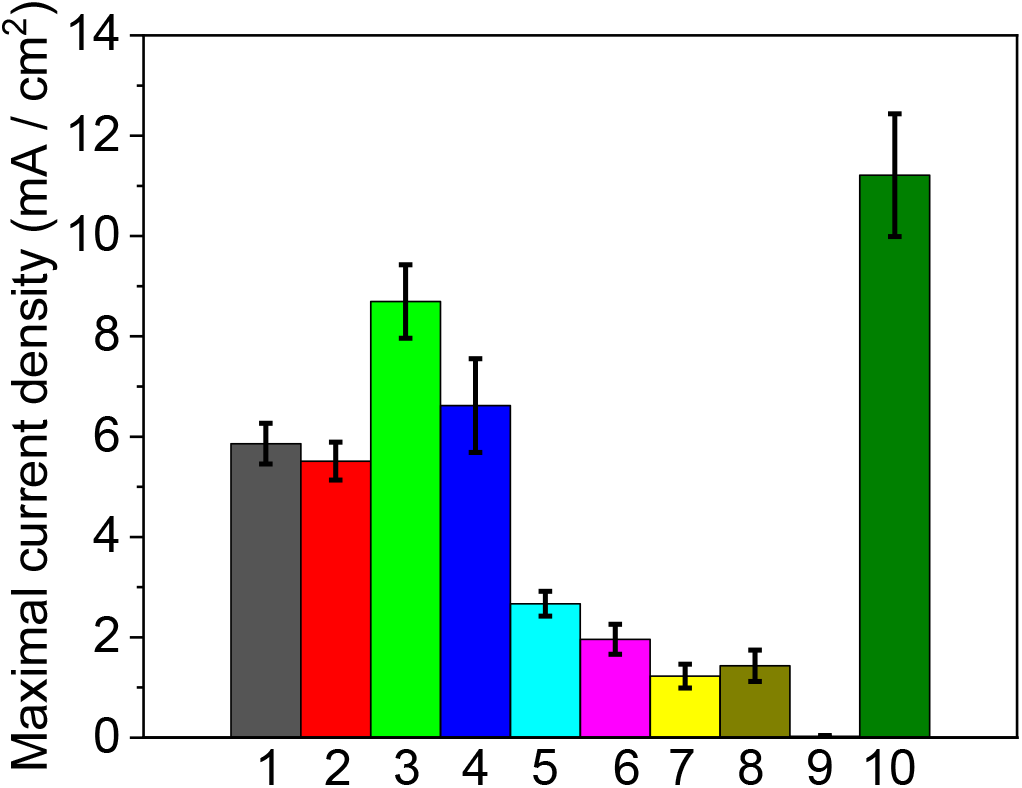
Photocurrent production of different plants. CA of leaves and the stems of ten different plants were determined. Maximal current densities production of Sage 5.86 +/- 0.41 mA/cm^2^ (1), Origanum 5.51 +/- 0.38 mA/cm^2^ (2), Moss 8.69 +/- 0.73 mA/cm^2^ (3), Cistus 6.62 +/- 0.93 mA/cm^2^ (4), Pine 2.67 +/- 0.25 mA/cm^2^ (5), Grapevine 1.96 +/- 0.3 mA/cm^2^ (6), Rose stem 1.22 +/- 0.23 mA/cm^2^ (7) Banana 1.43 +/- 0.31 mA/cm^2^ (8), *Opuntia ficus-indica* 0.03 +/- 0.02 mA/cm^2^ (9) and *Opuntia ficus-indica* after removal of its cuticle 11.21 +/- 1.22 mA/cm^2^ (10). The error bars represent the standard deviation over 3 independent measurements.

### 3.4 Photocurrent as a function of leaf development stage

In addition to outer surface permeability, we expected that the level of produced current in the BPEC would be proportional to the relative concentration of photosynthetic complexes in the leaf which change during leaf development (and/or senescence). To ascertain the correlation between leaf content and current, we examined *Cercis siliquastrum* leaves with different amounts of chlorophyll per square cm of leaf surface. In Fig. 4 we show the results of simultaneous chlorophyll. concentration, CA and the photo-increase of dissolved oxygen (ΔDO) measurements on 1 cm round segments of the different leaves (see Methods section for more details). The results show good correlation between the harvested photocurrents, the chlorophyll concentration, and the ΔDO values (which indicate the magnitude of photosynthetic activity from the entire illuminated leaf area). It should be reiterated that the kinetics of photosynthetic oxygen evolution and the reduction of the anode are different. As we have previously shown in our previous work (Shlosberg et al., 2021b), oxygen evolution initiates and terminates in a timescale of seconds when the light is turned on or off. Meanwhile, the transport of potential electron mediators (including NADPH) through the chloroplast, cytoplasm and cell membrane are slower. Therefore, upon illumination, the electron mediators accumulate in the cells before being secreted by the cells to reduce the anode. The chlorophyll *a/b* ratio in all three leaf types was ∼ 3.5, indicating that the pigment intensity is related to the concentration of photosynthetic units and not to changes in the composition of the complexes (with respect to the ratio of reaction centres to antenna proteins). These results also support our determination that photosynthesis is the source of current generation in the BPEC.

**Fig. 4.**
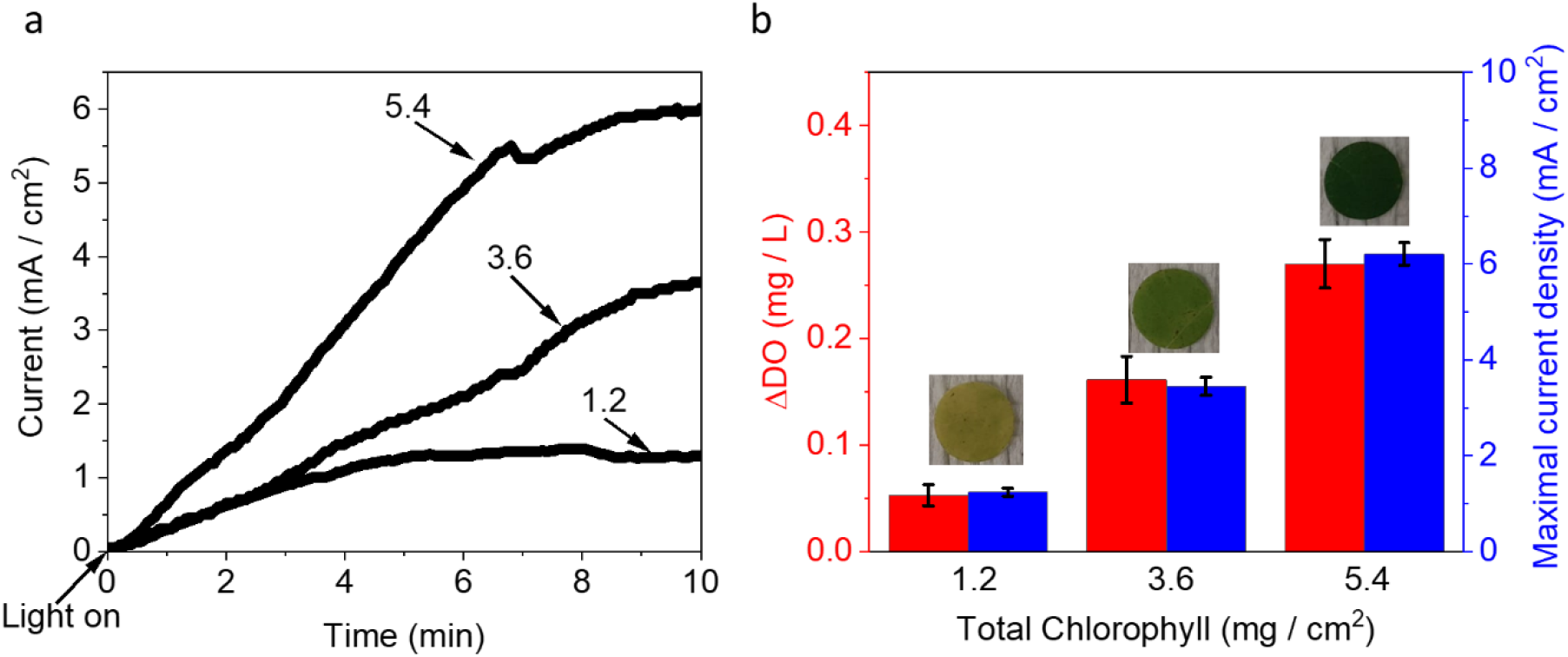
Photocurrent as a function of photosynthetic content. **a**. a typical CA measurement of leaves from *Cercis siliquastrum* at different developmental stages, containing different chlorophyll concentration levels, as indicated for each measurement. The numbers above each line represent total chlorophyll of the measured leaf (mg / cm^2^) **b**. Average measurements of the maximal current (blue) and dissolved oxygen (ΔDO, red) as a function of leaf total chlorophyll concentrations. leaves with total chlorophyll of 1.2, 3.6, and 5.4 mg/cm^2^ produced of 0.05 +/- 0.01, 0.16 +/- 0.02, and 0.27 +/- 0.02 mg/L O_2_, and maximal photocurrents of 1.23 +/- 0.08, 3.44 +/- 0.19, and 6.21 +/- 0.24 mA/cm^2^ respectively. Pictures of the leaves are shown above the bars. The error bars represent the standard deviation over 3 independent measurements.

**Fig. 5.**
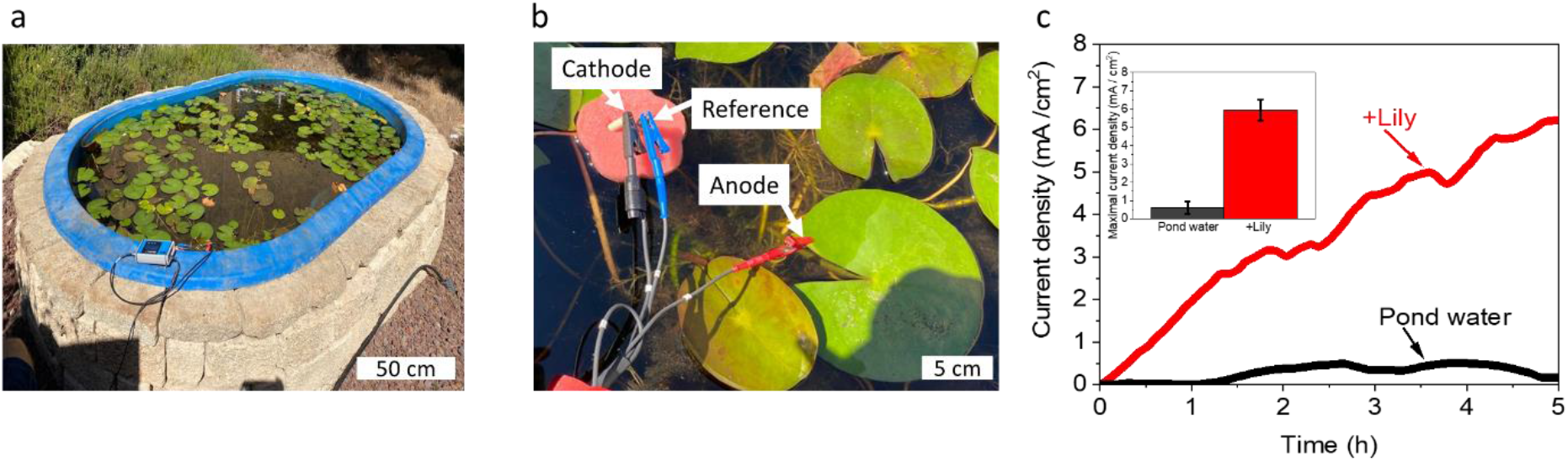
Towards applicative renewable energy technologies. CA of a Lily leaf was measured directly from its native growth environment using the water in the Lily Pond as the electrolyte of the BPEC. **a**. Photo of the Lily-pond with the measurement setup. **b**. An enlargement of the active area of the pond where the CA measurements were conducted. A stainless-steel clip anode grasps the Lily leaf. The cathode and reference electrodes are held by a sponge that floats on the water surface. **c** CA measurements in the Lily Pond water connected (red) or floating in the pond water (black) between the anode and the leaf. the maximal current produced in without and with a connection between the anode and the leaf were 0.003 +/- 0.002 and 0.017 +/- 0.005 mA / cm^2^ respevtively. The insert displays the maximal obtained currents over a 5-hr measurement. The error bars represent the standard deviation over 3 independent measurements.

### 3.5 Photocurrent production from Water Lilies in their native environment

In order to produce electricity from the leaves of trees or bushes (excluding succulents, see above), the leaves must be inserted into an aqueous solution. Unlike terrestrial plants, the native habitat of water plants can serve as an electrolyte, therefore may enable direct electricity generation without harming the plants. To demonstrate the potential of utilization of water plant based BPECs, we examined the ability of Water Lilies (*Nymphaeaceae*) in their native pond to generate photocurrents under the natural sunlight. CA measurements were conducted in the pond under the naturally changing sunlight intensity using an applied bias of 0.5 V. A stainless-steel anode clip was dipped in the pool held one of the leaves in the pond. A platinum wire cathode and an Ag/AgCl KCl 3M were held by a sponge, to float on the pond surface (Fig. 7a, b). Sunlight intensities of ∼ 200 – 900 μE / m^2^ /s and water temperatures of 13-17 ^0^C were measured during the duration of the CA measurement (Fig. S12). Current densities of 6.5 mA /cm^2^ was obtained in this configuration. If the anode is not associated with a leaf only a small current (∼0.5 mA/cm^2^) was obtained, most likely originating from corrosion or the existence of reducing molecules in the pond water. The pond water is of course a highly complex mixture of tap water, organic molecules, and a variety of insects and animals that inhabit the immediate environment.

### 3.6 A Model for electron transport in plant-based MET based BPECs

Based on the results we have presented here we propose a mechanism that explains the generation of photocurrents from intact leaves (Scheme. 1), We suggest that the main source of electrons originates from the natural photosynthetic linear electron pathway from water to NADPH. NADPH can then diffuse out the chloroplasts and the cell membrane to reduce the anode. (Shlosberg et al., 2021b, 2021c, 2020) The external electron transport occurs only under illumination and is inhibited by the addition of DCMU. In the case of plants with hard cuticles, electron transfer mediators are not freely released to the anode. Therefore, negligible currents are generated to the external anode or electrolyte solution. This can be overcome by removal of the external cuticle layer and rewiring of the inner softer photosynthetic tissue.

Based on the results presented here, the development of larger devices that utilize intact terrestrial plants to generate useful amounts of electrical currents and/or hydrogen. We show that while leaves with softer texture can generate higher photocurrents, currents can be generated from other photosynthetic tissues of almost any species. Our demonstration of the ability of water Lily ponds to generate photocurrents in their native habitat without the need to harvest them shows that by using the BPEC configuration described here, significantly more current can be obtained than previously reported for other aquatic plants (Hubenova and Mitov, 2015, 2012). As there is no need to process the leaves for generating photocurrents and the fact that almost any kind of leaves may be used, the economic cost of using plants as an electron source should be very low. Moreover, it may be integrated with crops that are grown for biodiesel production. Therefore, the ability of plants to directly generate electricity in BPECs without any processing holds great promise and potential for designing a “greener” future for energy technologies.

**Scheme. 1.**
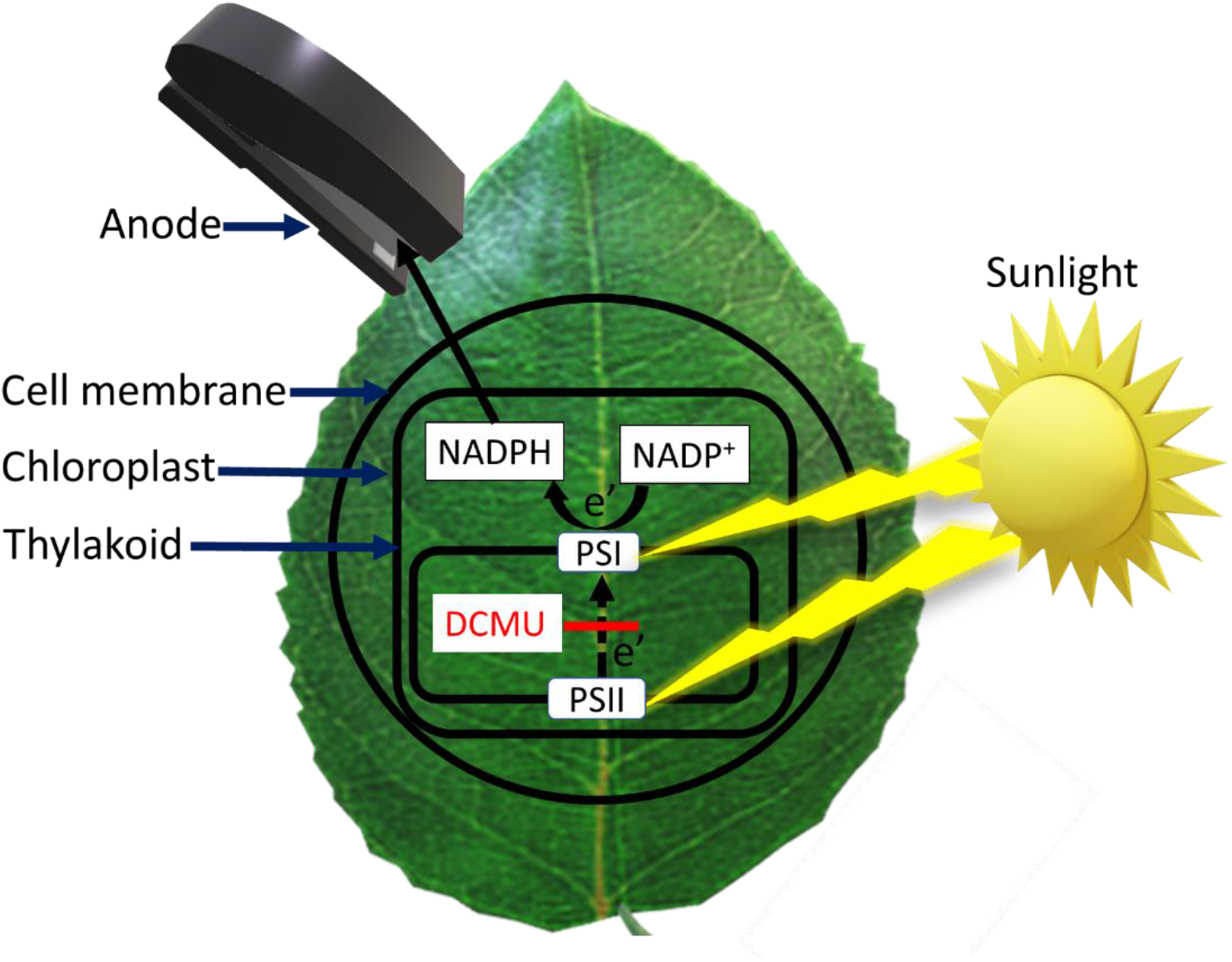
A model for the electron transport mechanisms in plant based BPECs. A Schematic model for the suggested electron transfer mechanisms in BPEC. **a**. The electron source is PSII which conducts water splitting under sunlight illumination. The electrons are being transferred by the multistep reduction reactions of the photosynthetic pathway through PSI to reduce NADP^+^ to NADPH. Under association with the electrochemical component of the BPEC, NADPH can exit the Chloroplast and the outer cell membrane to reduce the anode and produce current. The addition of DCMU blocks the electron transport from PSII and thus eliminates the photocurrent generation. Yellow stripes indicate the light illumination, dark black arrows mark the anode, cell membrane chloroplast, and thylakoid. Dashed arrows indicate electron transport. Whole black arrows indicate the trafficking of NADPH. A red line indicates the inhibition of the electron transport by DCMU.

## 4. Conclusions

In this study, we show the ability of intact higher plants (terrestrial and fresh water) to generate electrical currents in a novel bio-photoelectrochemical configuration. We further show that while permeable (soft surface) leaves can generate higher photocurrents than plants with hard outside surfaces, currents can be generated from other photosynthetic tissues of almost any species. We show that the electron source for the generated photocurrents derive from PSII as it is DCMU sensitive and utilize NADPH, released from the leaf, as one of the main electron mediating molecules. The concept of direct electricity production from plant leaves may be a base for the development of novel green energy technologies.

## Acknowledgements

Funding was provided by a “Nevet” grant from the Grand Technion Energy Program (GTEP) and a Technion VPR Berman Grant for Energy Research. Some of the results reported in this work were obtained using central facilities at the Technion’s Hydrogen Technologies Research Laboratory (HTRL) supported by the Nancy & Stephen Grand Technion Energy Program (GTEP), the ADELIS Foundation and the Solar Fuels I-CORE. We thank Dr. Rachel Edrei and Dr. Yifat Nakibly for technical support. Yaniv Shlosberg is supported by fellowships of the Nancy & Stephen Grand Technion Energy Program (GTEP) and by a Schulich Graduate fellowship.

## Author contributions

YS and NA conceived the idea. YS, OY and NA designed the experiments. YS performed the main experiments. MM assisted in performing parts of different experiments. YS, OY, GS and NA wrote the paper. NA and GS supervise the entire research project and provide funding.

## Competing interests

The authors declare no competing interests.

## Supplementary Information for

**Fig. S1.**
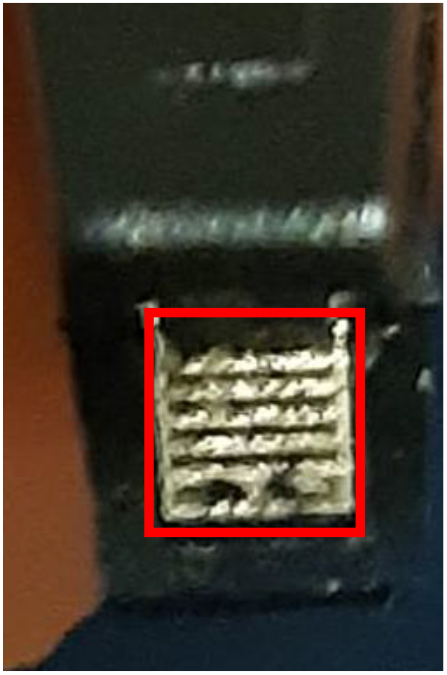
A picture of the edge of the clips that applies as anode. The inner stainless steel anode (Type 304) has a rectangular shape with non-sharp metallic stripes. A red rectangle shows the lower half of the anode that clasps the leaf.

**Fig. S2.**
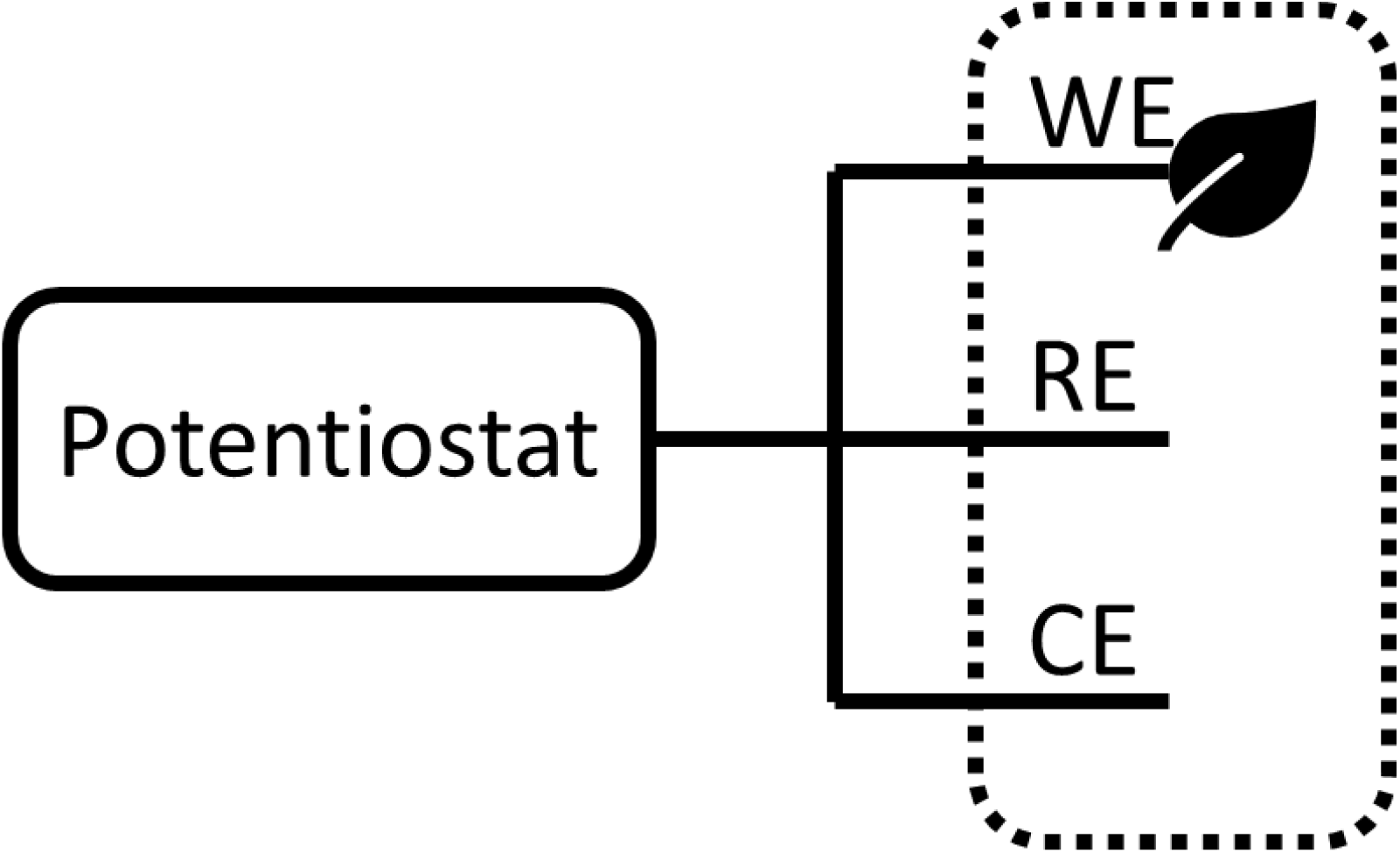
An electrical circuit diagram of the BPEC. WE, RE and CE represent the working reference and counter electrodes respectively. The leaf icon represent the grasp of the leaf by the WE. A dashed line represent the glass container with the NaCl electrolyte.

**Fig. S3.**
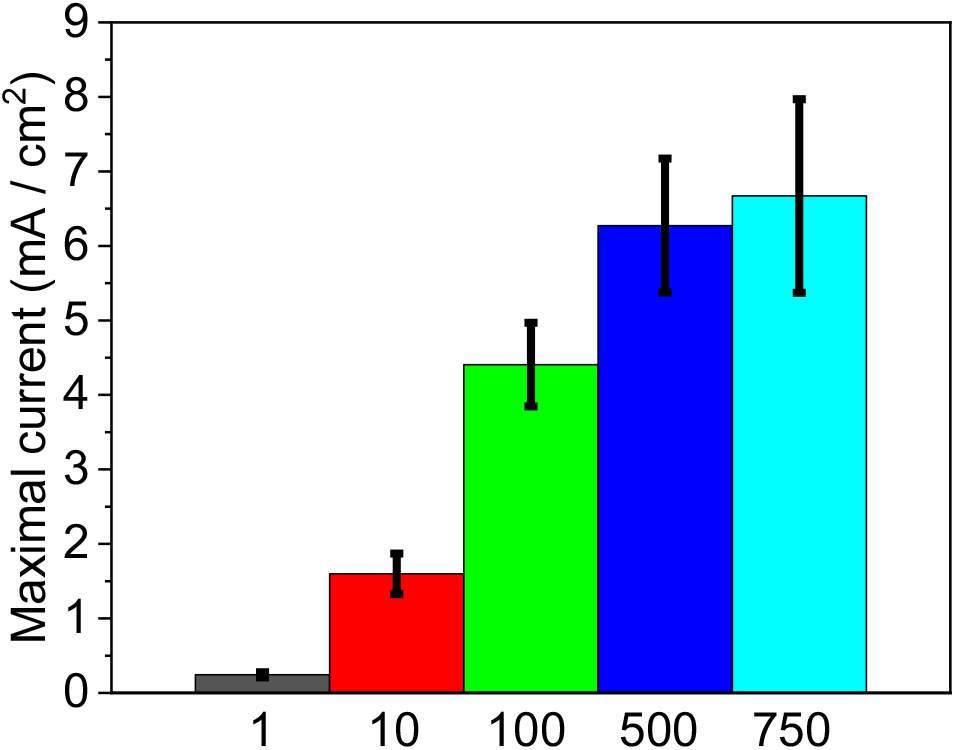
Determination of the optimal salinity for photocurrent production from spinach. CA of excised round pieces of spinach leaves with a diameter of 1 cm were measured for 10 min under solar irradiation (1 SUN) with an applied bias of 0.5 V on the anode. The measurements were conducted using increasing NaCl concentrations: 1, 10, 100, 500, or 750 mM. The bars represent the maximal photocurrent obtained after 10 min. The error bars represent the standard deviation over 3 independent measurements.

**Fig. S4.**
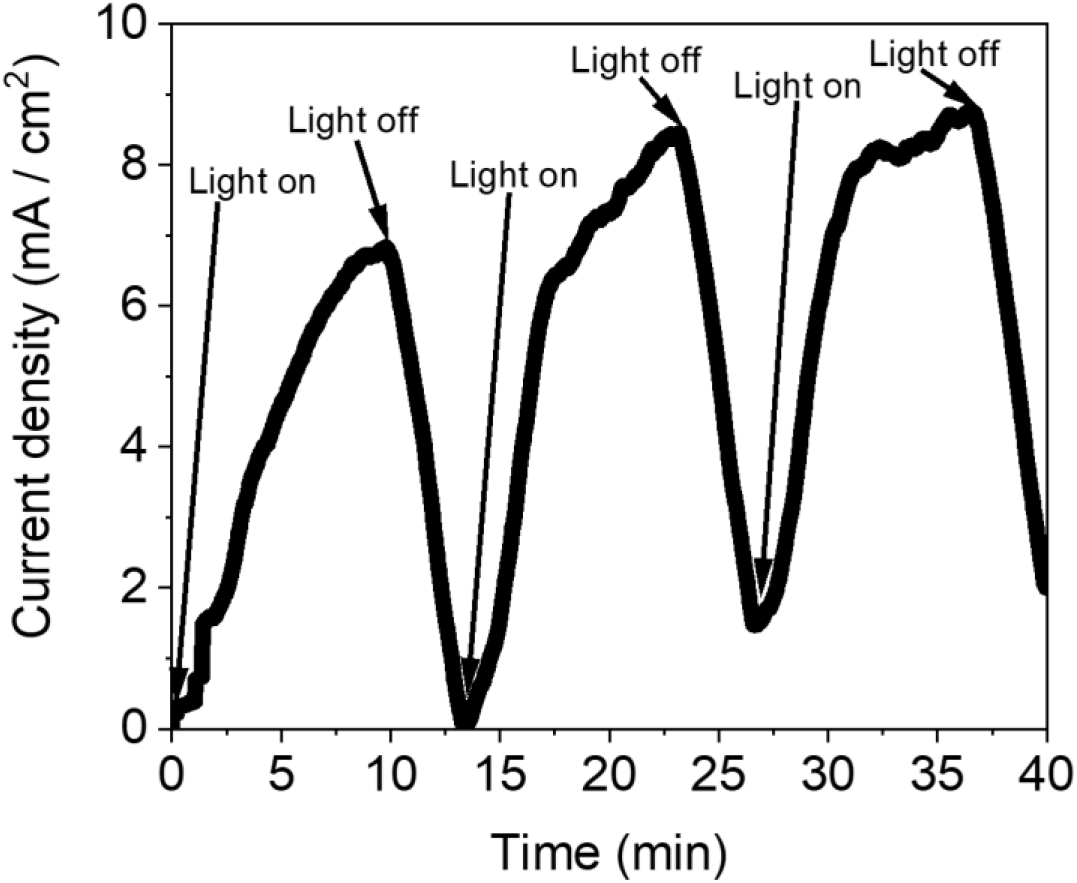
Current harvesting from leaves during light/dark intervals. CA of a spinach cutting was measured in intervals of 10 min. light/5 min. dark intervals. Initiation and termination of the illumination is marked by arrows.

**Fig. S5.**
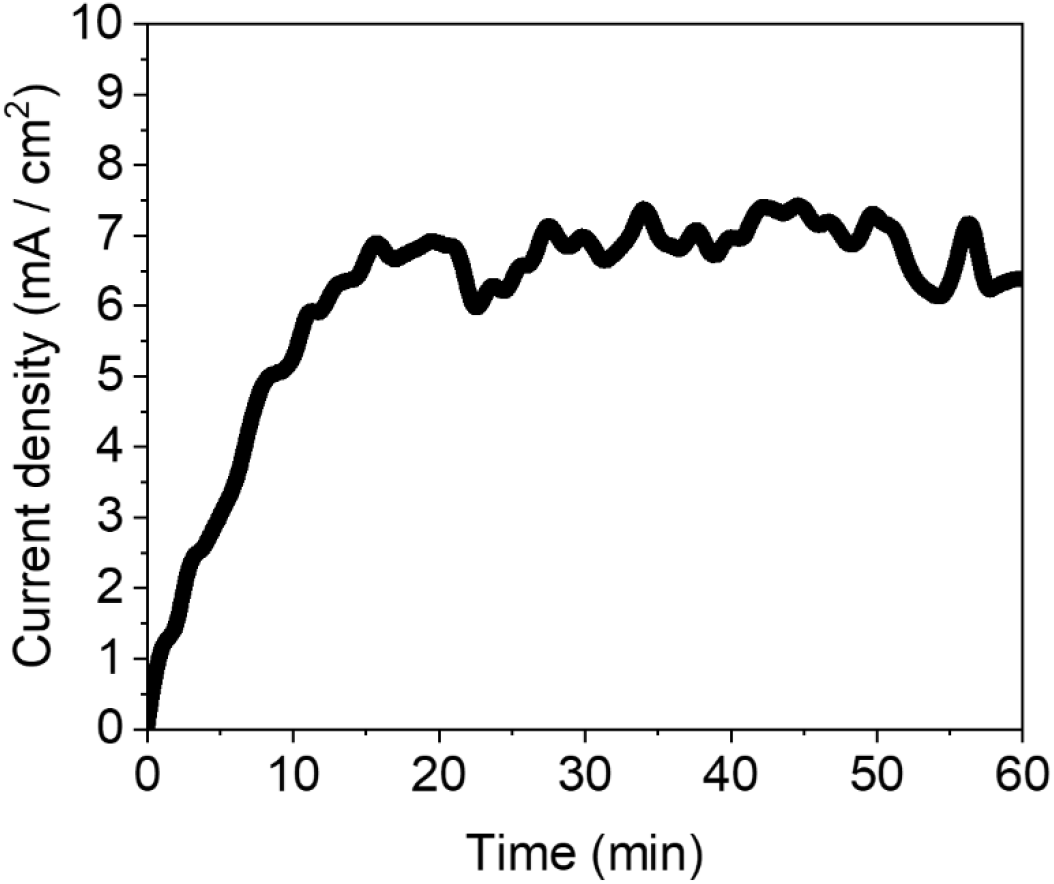
Evaluation of photocurrent production sustainability in a spinach based BPEC. CA of a spinach cutting was measured for 1 h.

**Fig. S6.**
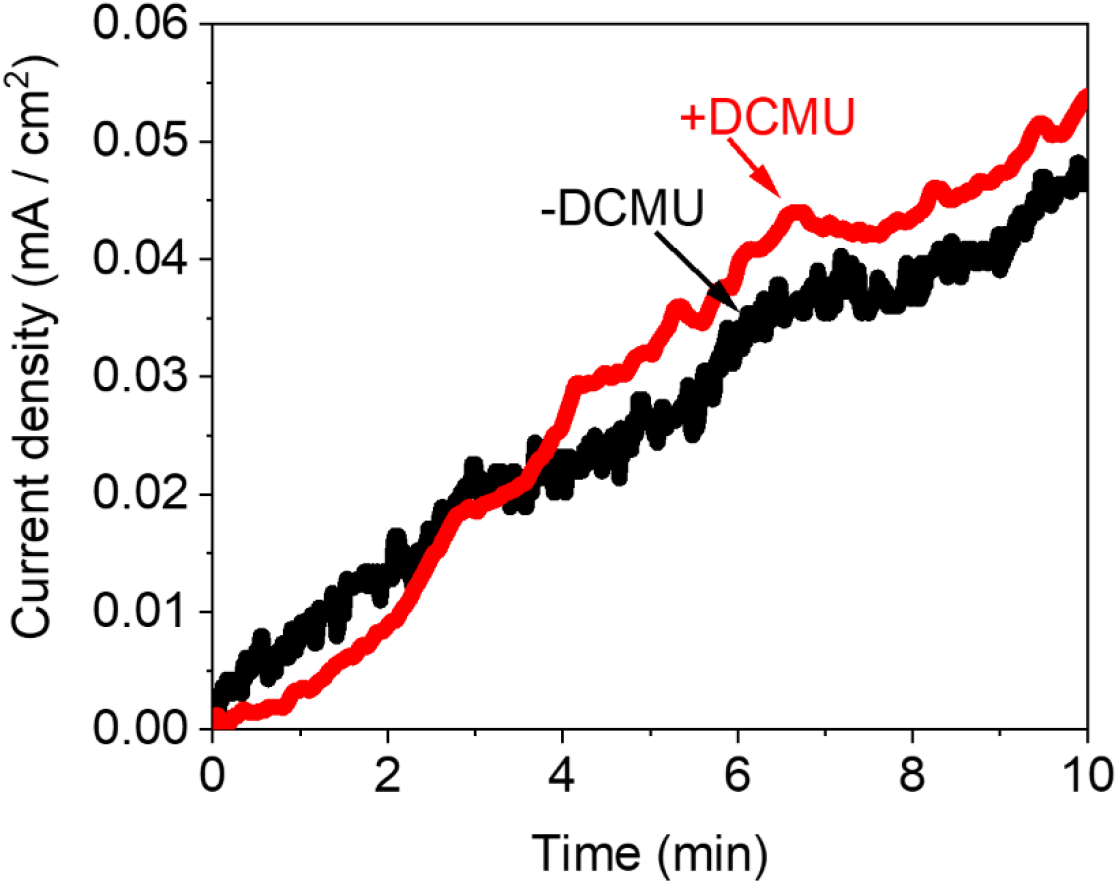
Estimation of the contribution of corrosion to the photocurrent. CA was measured with an empty stainless steel anode clips (without a leaf). The measurements were conducted in 0.5 M NaCl (black) or 0.5 M NaCl + 100 μM DCMU (red).

**Fig. S7.**
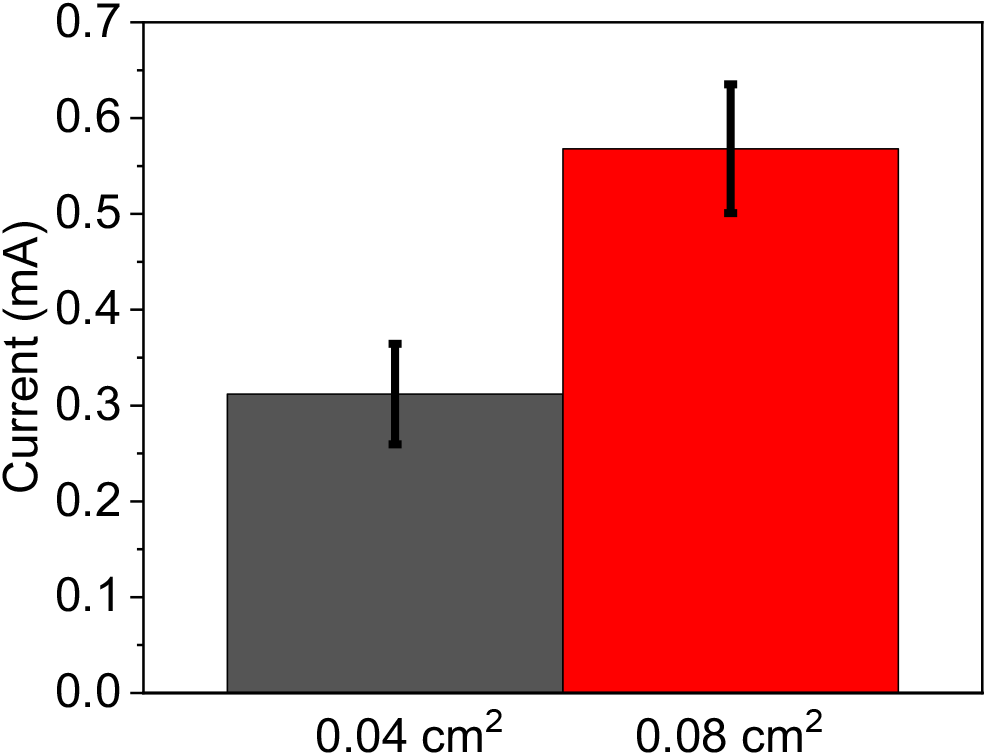
Photocurrent production using different anode sizes. CA of excised round spinach leaf pieces with a diameter of 1 cm were measured for 10 min under solar irradiation (1 SUN) with an applied bias of 0.5 V on the anode. The spinach pieces were directly touching both sides of the stainless-steel anode clips (anode area = 0.08 cm^2^), or just one side while the other side was tightly sealed with parafilm (anode area = 0.04 cm^2^). The bars represent the maximal photocurrent obtained after 10 min using an anode area of 0.04 cm^2^ (black), and 0.08 cm^2^ (red). The error bars represent the standard deviation over 3 independent measurements.

**Fig. S8.**
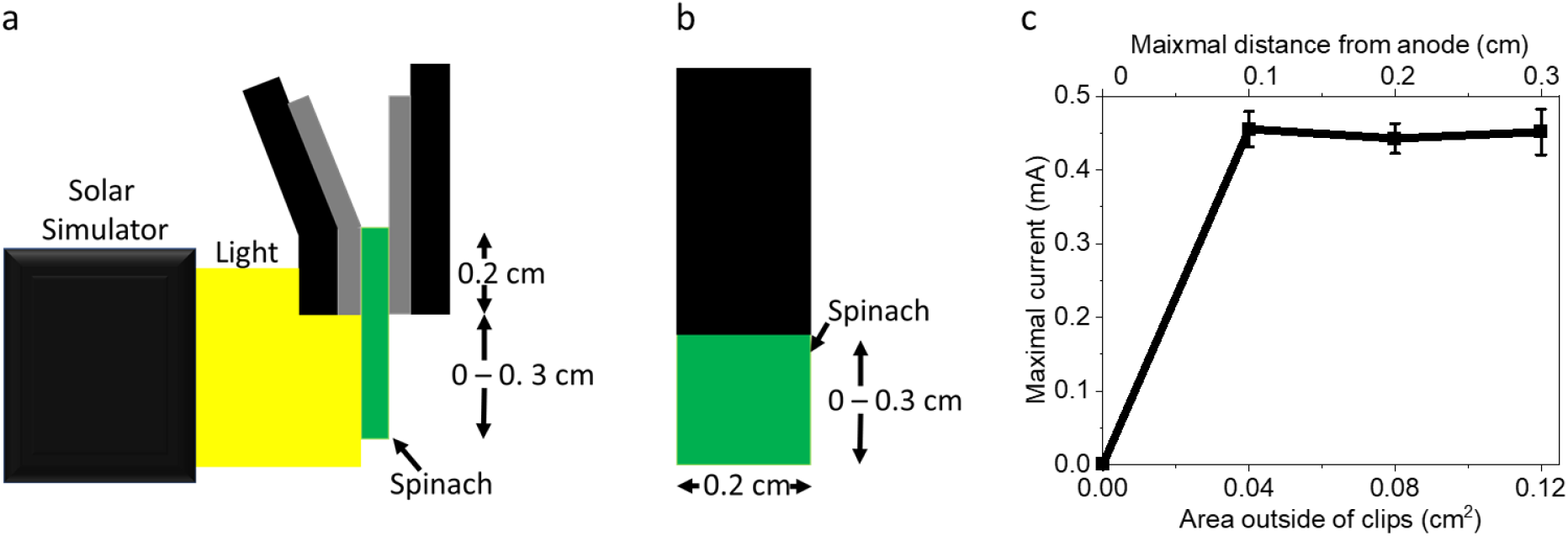
Evaluation of the maximal effective distance in the spinach leaf on the photocurrent. CA of excised spinach leaf pieces with fixed width of 0.2 cm and variable increasing lengths of 0.2 – 0.5 cm was measured for 10 min under solar irradiation (1 SUN). In all the measurements the upper part of the spinach tissue (0.2 cm) was fully covered by the anode clips. Schematic illustrations of the measurement setup, from the side (a) and front (b). Upon light irradiation, only areas of the of the spinach tissue outside of the clips were exposed to light. The fixed and variable dimensions of the anode clips and spinach piece are marked with black arrows. **c**. CA measurements of spinach pieces with different sizes. The Y-axis shows the maximal photocurrent at 10 min (without normalization to the anode area). The lower X-axis shows the area of the leaf that is not covered by the anode clips and exposed to light. The upper X-axis shows the maximal distance of the leaf from the anode. The error bars represent the standard deviation over 3 independent measurements.

**Fig. S9.**
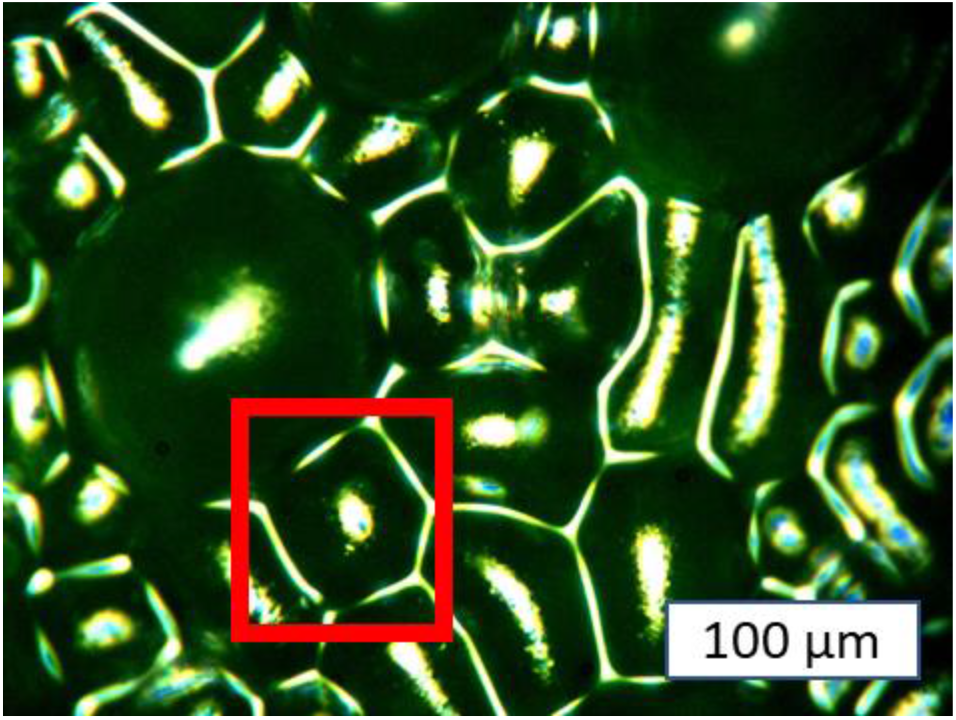
Microscopic observation of the grasped spinach cutting. A microscopic photo of the spinach cutting after 20 min incubation in 0.5M NaCl being grasped by the stainless-steel anode. A red rectangle shows intact cells wall boundaries of the representative cells (surrounding the large stomata) in the spinach tissue.

**Fig. S10.**
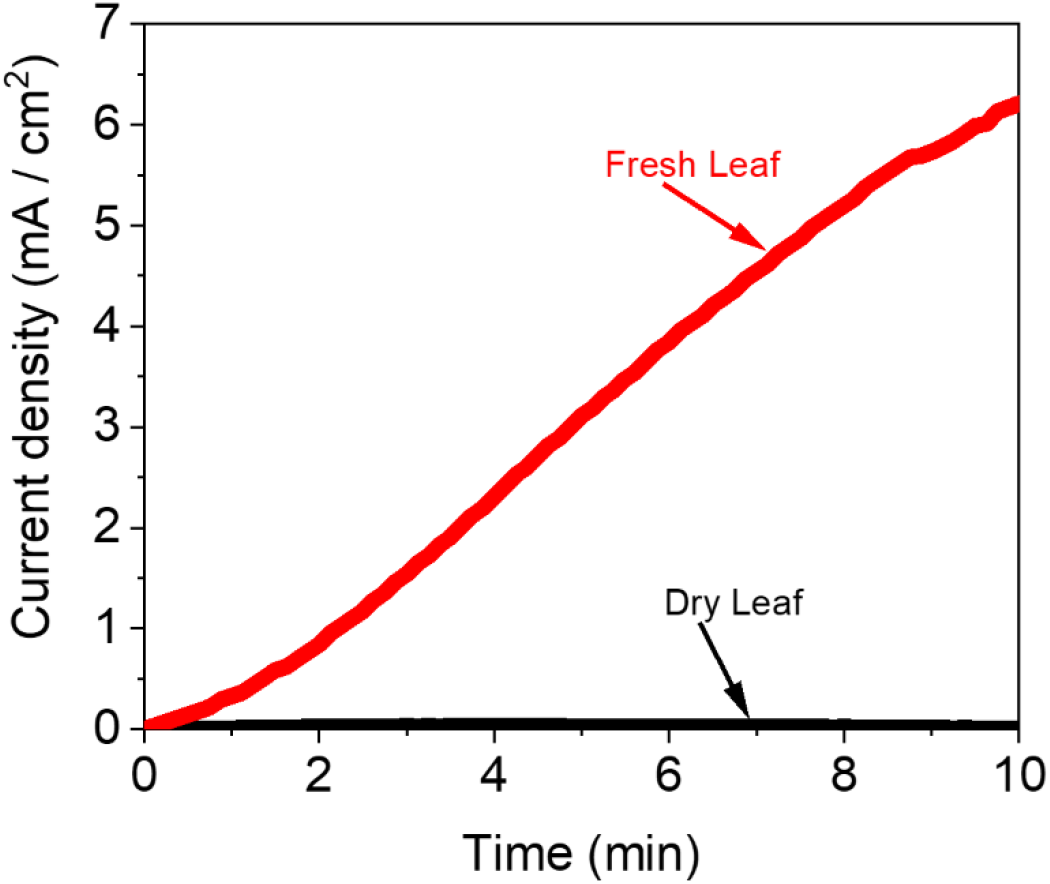
Dry leaves do not produce photocurrent. CA measurements of dry (black) and fresh (red) cistus leaves were conducted in light over 10 min.

**Fig. S11.**
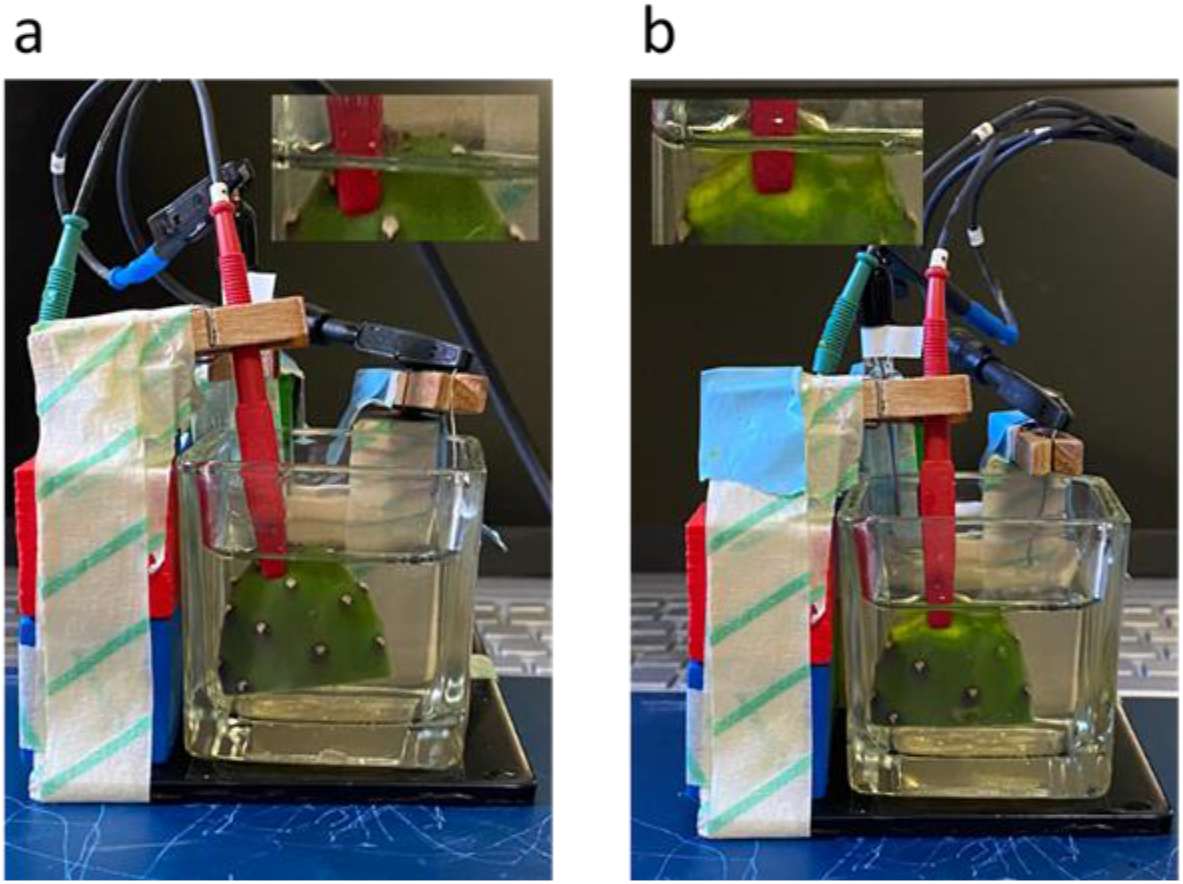
Pictures of the experimental setup for CA measurements of *Opuntia ficus-indica*. The measurements were conducted in three electrode mode using the stainless-steel clip as anode, a platinum wire as cathode, and Ag/AgCl 3M NaCl as a reference electrode, with an applied electric potential bias of 0.5 V on the anode in 0.5 M NaCl solution. Intact or peeled leave-like stems of *Opuntia ficus-indica* were held by the anode clips. A platinum cathode and Ag/AgCl 3M NaCl reference electrode float on the surface. **a**. Intact leaf. **b**. Peeled leaf. The inserts show an enlargement of the anode clips that holds the intact and a peeled leaf.

**Fig. S12.**
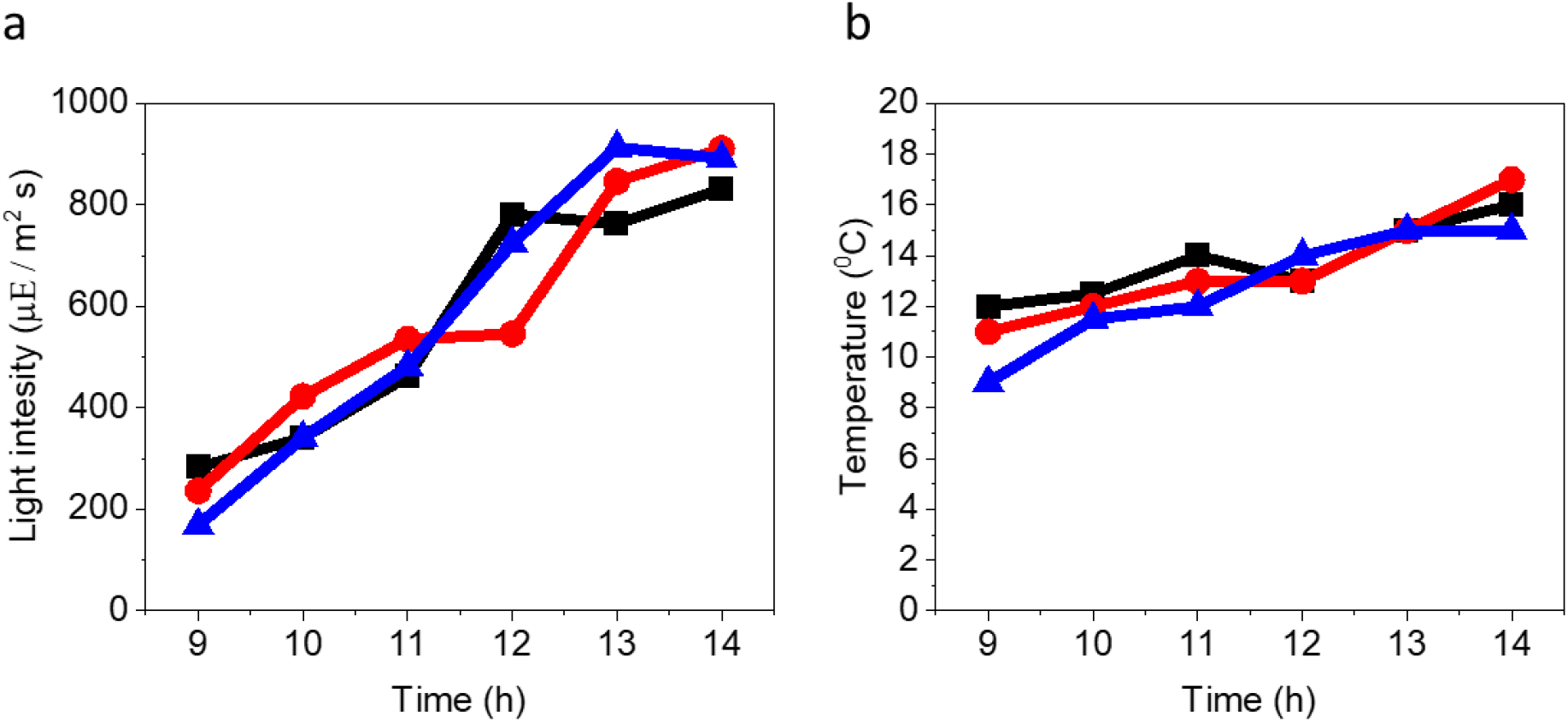
Temperature and light intensity measurements in the Lily Pond used for CA measurements. Temperature and light intensity were measured for 5 h (9:00 – 14:00) in parallel to the CA measurements in the Lily-pond. **a**. Light-intensity measurements. **b**. Temperature measurements. The black, red and blue lines represent 3 independent measurements. The numbers at the X-axis represent the time of the day when the measurements were conducted.

## Notes

### Competing Interest Statement

The authors have declared no competing interest.

### Summary of Updates

The manuscript was changed.

